# Assessment of Visual Function in Mice Using Light/Dark Box and Multi-Feature Machine Learning

**DOI:** 10.1101/2025.11.07.687135

**Authors:** Tengxiao Wang, Karen Chang, Matteo Tomasi, Chen-Yuan Lee, Dong Feng Chen, Gang Luo

## Abstract

The light/dark box test can be used to assess visual function in rodents based on their spontaneous behavior in response to light. Commonly used assay relies on a single behavioral metric, dwell time in the light or dark compartment, which may be influenced by factors other than vision, leading to unreliable assessment results. To overcome this, we developed a multi-feature machine learning paradigm by extracting multiple mouse behavioral metrics, standardizing them as features to train machine learning models, thereby achieving reliable and automated vision assessment. We systematically compared the classification performance of single-metric versus multi-feature machine learning approaches in sighted and blind mice, using wild-type and rhodopsin-deficient mice, with a subset further subjected to double optic nerve crush. We found that the multi-feature method can improve classification performance and exhibit great robustness to different experimental settings. Additionally, we further improved model performance by applying feature importance analysis and constructing an optimized feature subset. These findings suggest that the reliability of commonly used single dwell time measure for vision assessment could become unreliable, as shown in our experiment, probably because in addition to vision other factors also impact dwell time. Our study demonstrated an improved assessment method based on a combination of multiple behavior features through machine learning.

**Author Summary:** Assessing visual function in mice is essential for studying eye diseases and drug development. The light/dark box test evaluates visual function by measuring the spontaneous behavioral response of mice to light, providing a training-free behavioral approach that helps simplify the assessment process and improve research efficiency. However, traditional light/dark box tests rely on a single behavioral metric, dwell time in the light or dark compartment, to assess visual function, which may be influenced by factors other than vision, such as anxiety and exploratory behavior, leading to limited reliability of assessment results. Here, we demonstrate that integrating multiple behavioral features through machine learning can improve the reliability and stability of vision assessment. By automatically tracking and analyzing various behavioral metrics of mice, such as movement patterns, speed, and spatial preferences, the proposed method can more reliably distinguish between sighted and blind mice. Furthermore, the method demonstrates stable performance across different experimental settings, showing good applicability. This automated, reliable, and easily generalizable method can provide a convenient and efficient means for visual assessment in preclinical research, facilitating vision disease research and drug development.

## Introduction

Vision is one of the most critical sensory modalities for rodents, playing a vital role in navigation, foraging, and threat avoidance. In preclinical research, mice are widely used as models to investigate genetic and acquired visual disorders, making accurate assessment of their visual function essential. Many studies have primarily relied on behavioral training paradigms^[1]-[4]^ or electrophysiological recordings^[5]-[7]^ to assess visual capabilities in mice. Behavioral methods typically rely on constructing visual mazes or similar apparatuses, where mice must undergo repeated training to associate visual stimuli with specific responses. These procedures are time-consuming and not easily scalable. Electrophysiological approaches, such as recording visual evoked potentials, assess visual function by measuring neural responses to visual stimulation, but often involve invasive procedures that limit their suitability for long-term tracking of visual development or functional recovery following interventions^[2]^. As a result, there is increasing needs for noninvasive, and training-free approaches that enable rapid and reliable assessment of visual function in mice, especially across different strains, ages, sexes and pathologies.

The light/dark box is a classical behavioral apparatus originally developed to assess anxiety levels in rodents^[8]^, and has been also adopted for assessing visual function^[9]-[24]^. The setup typically consists of a light compartment and a dark compartment connected by an opening. During the experiment, mice are allowed to move freely between the two areas. Their behavior is believed to reflect an innate response to light stimulation. The assumption is that mice with at least some level of vision typically exhibit aversion to the illuminated area, while blind mice may show no clear preference or a delayed response. Thus, the light/dark box offers a training-free method for assessing light perception in mice.

Most existing studies assess visual function in mice by quantifying the dwell time in the light or dark compartments of the light/dark box^[9]-[22]^. In addition, some other behavior measures have also been explored. Lagali *et al*.^[23]^ evaluated visual responsiveness by measuring changes in speed before and after light exposure. Shindo *et al*.^[24]^ compared multiple behavioral metrics separately, including dwell time in the light compartment, time to enter the dark compartment, and the number of transitions between compartments, each assessed independently. While these assessments based on individual metrics have demonstrated some ability to differentiate between sighted and blind mice, they have several limitations. In previously studies, impacts from confounding factors such as individual variability, anxiety levels, or exploratory motivation have been reported^[8]^. These confounding factors can lead to unreliable assessment and increase the likelihood of misclassification. In addition, there are also a considerale variation in experimental protocols across studies. A typical assessment procedure involves placing the mouse in the apparatus for a brief habituation period, followed by a predefined observation window during which behavioral metrics are recorded. According to literature, the duration of habituation ranges from 1-5 minutes, and the observation window varies from 3-10 minutes^[9]-[16], [18]-[24]^. Whether such differences in procedural settings influence the experimental outcome remains unclear, and this is also a question that the present study aims to investigate.

In this paper, we propose a novel machine learning paradigm based on multiple behavioral features to enhance the accuracy and generalizability of visual function assessment in the light/dark box system. Through feature importance analysis, we further investigated approaches to enhance the model’s performance. This unified framework enables automated evaluation of visual performance in mice, with the potential to accelerate vision-related drug discovery and support large-scale behavioral phenotyping in preclinical studies.

## Results

### Enhanced discriminative power of multi-feature assessment versus dwell time alone

As shown by the percentage of dwell time in the dark compartment (Fig. 1A, B) and the machine learning (ML) model-predicted probabilities (Fig. 1C, D), we first compared the discriminative ability of single-metric versus multi-feature assessments under different observation window settings. Fig. 1A illustrates the effect of window length. The starting time of the observation window is fixed at 0 and the observation window length extends from 1 to 10 minutes. Fig. 1B shows the results under different duration of habituation settings. Here, we fixed the window length at 5 minutes (the most common experimental setting^[9]-[13], [16], [18]-[19], [21]-[22], [24]^) and shifted the window start time from 0 to 10 minutes, thereby altering the duration of habituation. The results in Fig. 1A and B demonstrate that the dwell time distributions of the sighted and blind groups were highly overlapped, with no statistically significant differences observed under any experimental setting (smallest *p* = 0.252 at 2-min window length in Fig. 1A; smallest *p* = 0.129 at 3-min start time in Fig. 1B; t-test). This indicates that single dwell-time assessment provided limited discriminative power under our experimental settings. It is also shown in Fig. 1A and B that the blind group showed larger box heights and whisker spans, indicating greater individual variability in dark-side dwell time compared with sighted mice. This finding indicates that in the absence of visual drive, mouse behavior may be more readily influenced by non-visual factors, resulting in greater dispersion of the measure and further reducing the reliability of single dwell-time as a discriminative metric.

**Fig. 1.**
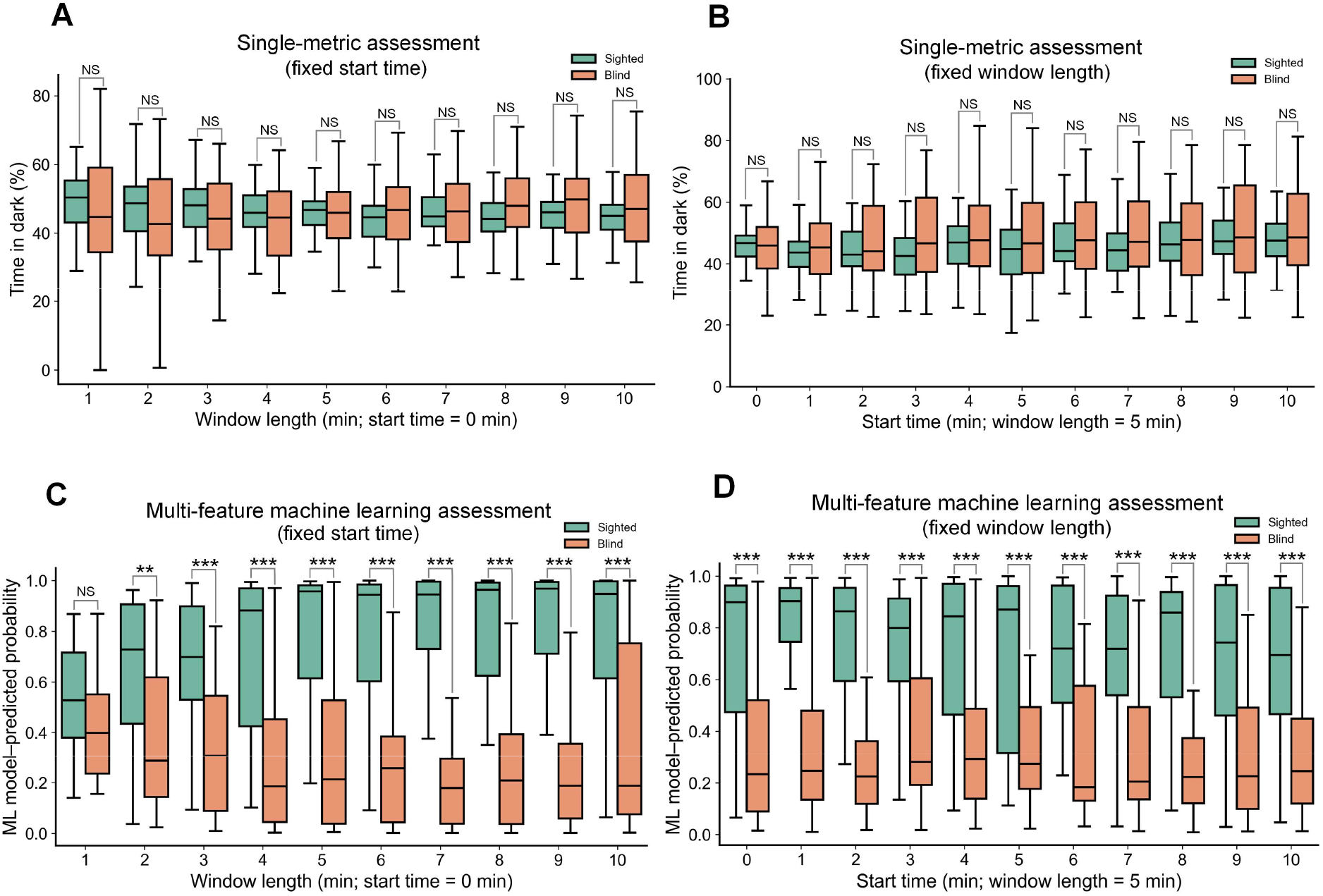
Comparison of single-metric and multi-feature machine learning approaches for visual function assessment in mice. **A**. Single-meatric assessment using dark dwell time percentage with observation window lengths of 1-10 minutes and start time fixed at 0 minutes. **B**. Single-meatric assessment with observation window length fixed at 5 minutes and start times of 0-10 minutes. **C**. Multi-feature machine learning (ML) assessment with observation window lengths of 1-10 minutes and start time fixed at 0 minutes. **D**. Multi-feature ML assessment with observation window length fixed at 5 minutes and start times of 0-10 minutes. Green box plots represent sighted mice (*n*=25), orange box plots represent blind mice (*n*=25). Box plots show median, interquartile range, and whiskers extending to 1.5× interquartile range. Statistical significance was determined by t-test. NS = not significant (*p* ≥ 0.05); \*\**p* < 0.01; \*\*\**p* < 0.001.

Fig. 1C and D illustrate the discriminative ability of the ML approach based on multiple behavioral features. Specifically, we employed a support vector machine (SVM) model, using ten behavioral metrics extracted from each observation window as input features, and adopted the model-predicted class probabilities as a replacement for the single dwell-time assessment. In Fig. 1C, with the start time of the observation window fixed at 0, increasing the window length from 1 to 10 minutes led to progressively greater separation between sighted and blind groups. At the 1-minute window, no significant difference was observed between the two groups (*p* = 0.155), whereas significant group differences emerged starting from 2 minutes (*p* = 0.002 at 2 minutes; *p* < 0.001 for window lengths greater than 2 minutes), with this separation remaining stable across longer observation windows. In Fig. 1D, when the window length was fixed at 5 minutes and the start time was shifted from 0 to 10 minutes, significant differences between the two groups were consistently observed under all conditions (*p* < 0.001). The results of Fig. 1C and D indicate that when the observation window reaches adequate duration (2 minutes or longer in this study), the multi-feature machine learning method can significantly discriminate between sighted and blind mice, and is insensitive to the duration of habituation. Overall, compared with single-metric assessment, the multi-feature machine learning approach provides substantially improved discriminative ability for visual function, is less affected by experimental settings, and exhibits greater reliability and robustness. Details of model training and feature definitions are provided in the Methods section.

### Enhanced classification performance of multi-feature assessment across observation time windows

We further compared the classification performance of single-behavior meatric assessment and multi-behavior feature ML assessment across observation window lengths ranging from 1 to 10 minutes. Instead of fixing the window start time, we implemented a sliding window approach where windows of each length moved along the temporal axis at 0.5-minute intervals, thereby fully utilizing temporal information across the entire 20-minute experimental session and providing a more comprehensive evaluation of model performance across different time scales. As shown in Fig. 2A, we first employed the receiver operating characteristic (ROC) curve to assess model performance. The area under the curve (AUC) is a standard metric quantifying the overall discriminative ability of a classifier, ranging from 0.5 to 1.0, with values closer to 1 indicating stronger classification ability, and AUC = 0.5 reflecting random-level classification. The results showed that the multi-feature assessment consistently outperformed the single-metric assessment across all window lengths. The AUC of the multi-feature model increased with window length, reaching a peak of 0.836 at 7 and 8 minutes, and remained high at 9 and 10 minutes (0.823 and 0.819, respectively). By contrast, the single-metric model showed only marginal improvement with increasing window length, but its AUC never exceeded 0.60, indicating performance only slightly above random chance. Taking the 7-minute window as an example, we plotted the ROC curves of both models. The AUC of the multi-feature assessment was 0.836, compared to only 0.552 for the single-metric assessment. We also highlighted the optimal threshold on each ROC curve, defined as the point with the maximum sum of sensitivity and specificity. For the multi-feature model, the optimal threshold was 0.519, meaning that each sample was classified as sighted when the predicted probability exceeded 0.519 and as blind otherwise. Under this criterion, the sensitivity and specificity reached 0.741 and 0.833, respectively, indicating that the model correctly identified 74.1% of sighted mice and 83.3% of blind mice. In contrast, for the single-metric model, the optimal threshold corresponded to 59.75% dark dwell time, at which the sensitivity was relatively high (0.881) but the specificity was extremely low (0.267), resulting in a strong bias toward false positives and overall poor classification performance.

**Fig. 2.**
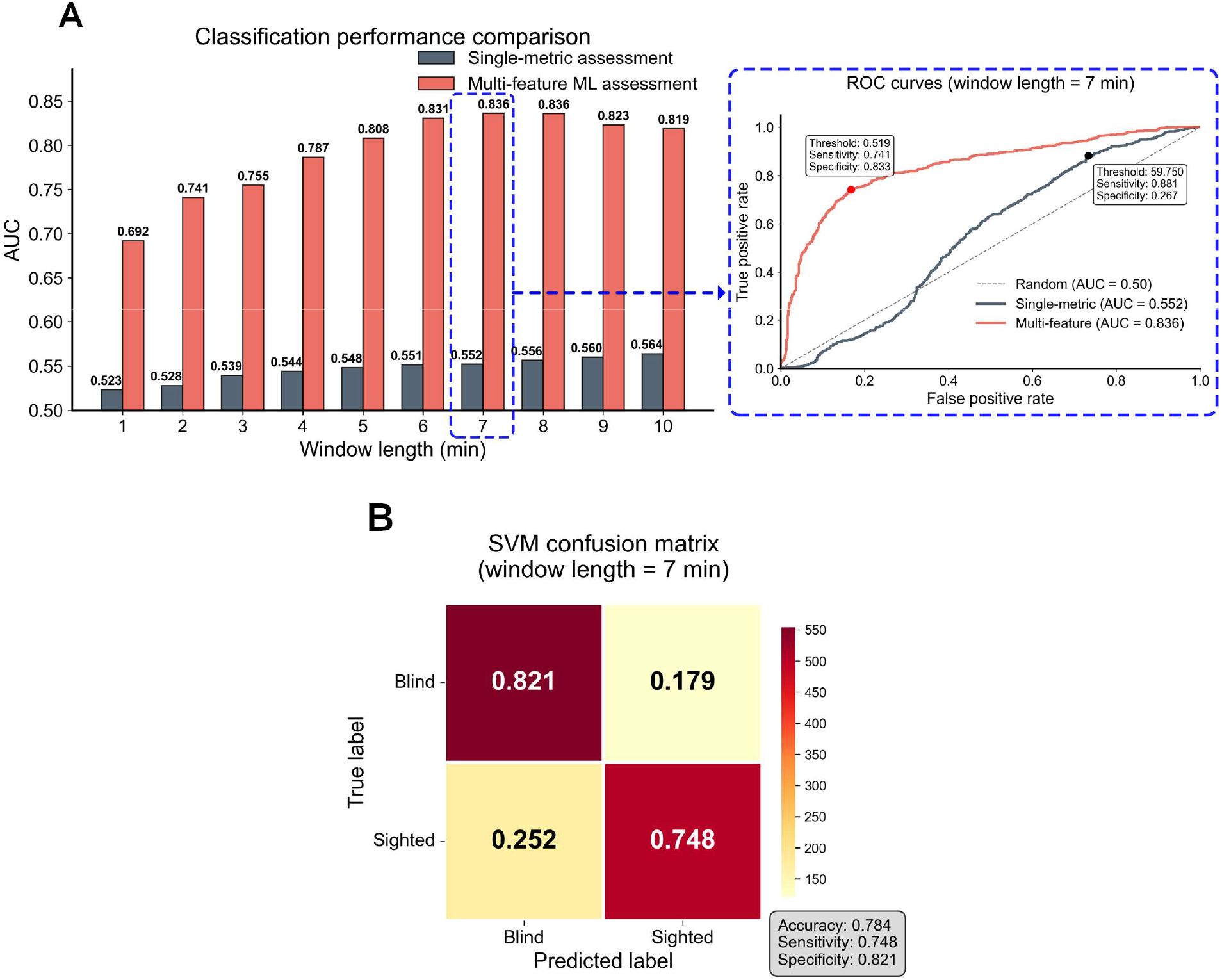
Comparison of classification performance across different observation window lengths. **A**. Receiver operating characteristic (ROC) analysis of single-metric and multi-feature models under observation window lengths of 1-10 minutes, with each window slid forward at 0.5-minute intervals. Orange line represents the multi-feature model (AUC = 0.836 at 7 min), and gray line represents the single-metric model (AUC = 0.552 at 7 min). Insets show the ROC curves at the 7-minute window, with the optimal thresholds indicated. **B**. Confusion matrix of the multi-feature model at a 7-minute window using the default threshold of 0.5. The model achieved an overall accuracy of 0.784, with sensitivity of 0.748 and specificity of 0.821.

As shown in Fig. 2B, we further employed a confusion matrix to illustrate the detailed classification outcomes of the multi-feature model across different categories, while also evaluating its accuracy. It is important to note that AUC and accuracy are conceptually different: AUC is derived from the ROC curve and reflects the model’s overall classification ability across all possible thresholds, whereas accuracy depends on predictions at a fixed threshold and directly reflects the model’s performance under that setting. Consequently, AUC values are generally higher than single-point accuracy. When constructing the confusion matrix, we adopted the default threshold of 0.5, which is the most common setting for classifiers, rather than the optimal threshold of 0.519 identified from the ROC curve. In this analysis, a total of 1350 samples were used under the 7-minute observation window, comprising 675 sighted and 675 blind mice samples. Among the blind samples, 554 (0.821) were correctly classified as blind, while 121 (0.179) were misclassified as sighted. Among the sighted samples, 505 (0.748) were correctly identified, whereas 170 (0.252) were misclassified as blind. Overall, the model achieved a classification accuracy of 0.784, with a sensitivity of 0.748 and a specificity of 0.821.

### Comparative analysis of machine learning classifiers

Fig. 3 compares the inference performance of five commonly used machine learning classifiers across observation window lengths of 1-10 minutes, including support vector machine (SVM), logistic regression (LR), multilayer perceptron (MLP), random forest (RF), and extreme gradient boosting (XGBoost). These models include kernel-based learning (SVM), linear classification (LR), neural networks (MLP), and ensemble tree-based approaches (RF and XGBoost). They are well suited for the present study with a relatively limited dataset, providing a reasonable balance between model complexity and generalization capability. The detailed training settings for each model are provided in the Methods section.

**Fig. 3.**
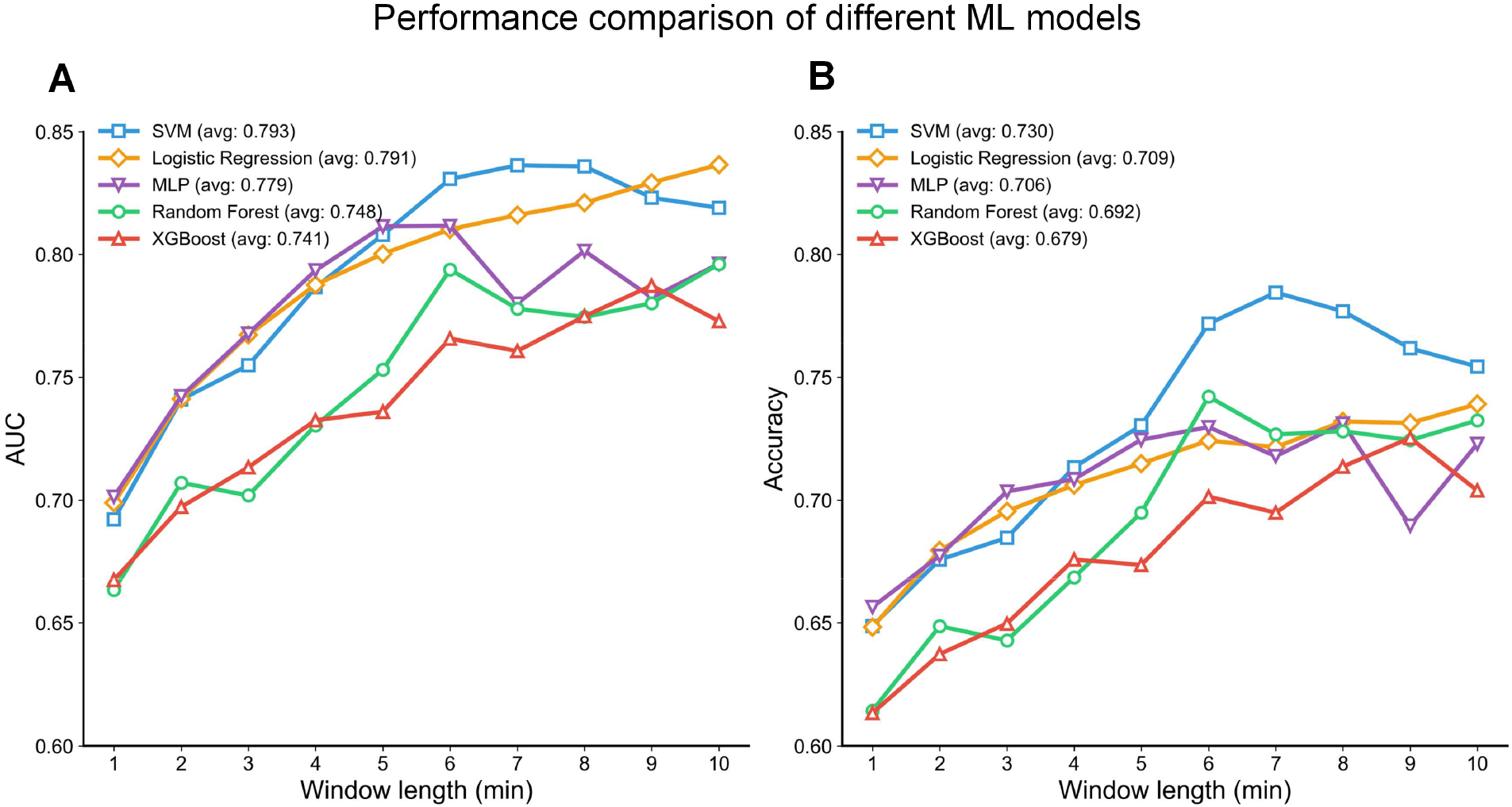
Comparison of machine learning models. **A**. AUC and **B**. classification accuracy of five classifiers across observation window lengths of 1-10 minutes. Models include support vector machine (SVM), logistic regression (LR), multilayer perceptron (MLP), random forest (RF), and extreme gradient boosting (XGBoost). SVM achieved the highest overall performance, followed by LR, whereas MLP exhibited intermediate results and RF and XGBoost showed relatively limited predictive ability. Average AUC and accuracy values across all window lengths are reported in the legends.

Performance analysis showed that the AUC and accuracy of all models gradually increased with longer observation windows and became relatively stable within the 6-10 minute range. Specifically, SVM achieved the best performance across most window lengths, with an average AUC of 0.793 and an average accuracy of 0.730, reaching its peak at 7 minutes before showing a slight decline. The LR model ranked second, with an average AUC of 0.791 and an average accuracy of 0.709, its AUC continued to improve at longer windows (≥ 9 minutes) and exceeded that of SVM, although its accuracy remained lower than SVM. The MLP model exhibited intermediate performance, with an average AUC of 0.779 and an average accuracy of 0.706. RF and XGBoost demonstrated relatively lower overall performance, with RF achieving an average AUC of 0.748 and an average accuracy of 0.692, and XGBoost achieving an average AUC of 0.741 and an average accuracy of 0.679. It is noteworthy that AUC values consistently exceeded accuracy scores across all models. This phenomenon can be attributed to the fundamental differences in these evaluation metrics: AUC measures the model’s overall discriminative capacity across all possible decision thresholds, whereas accuracy is dependent on a specific fixed threshold and is therefore more susceptible to the influence of class distribution and threshold selection. In other words, AUC better reflects the model’s inherent discriminative power, while accuracy represents the actual classification performance at a particular threshold. Overall, SVM achieved the highest performance, followed by LR, whereas MLP yielded intermediate results and RF and XGBoost demonstrated comparatively lower predictive performance.

### Feature importance analysis and subset optimization

Fig. 4 illustrates how feature importance analysis was used to identify an optimized feature subset that further improves classification performance. (A) We first applied the RF model to evaluate the importance of all 10 behavioral features listed in Fig. 4A and ranked them in descending order. Due to the impurity-based splitting mechanism inherent to RF, it is naturally capable of computing feature importance as part of its algorithmic structure, without requiring additional computation, which makes the results more interpretable. In Fig. 4A, the feature importance values are normalized such that the sum of all feature importances equals 1, allowing direct comparison across features. The analysis showed that “entries to dark (F1),” “dark average speed (F2),” and “light average speed (F3)” had the highest importance, each with a weight greater than 0.1, while the remaining features were less important. (B) To verify the impact of feature importance on classification performance, we conducted a stepwise feature ablation experiment (Fig. 4B) on the RF model, following the standard practice of progressively removing features from least to most important. Features were sequentially removed according to their inverse importance ranking, and the resulting changes in AUC and accuracy were assessed. Interestingly, we found that removing certain features (F9, F8, F7, F5) did not reduce but instead improved model performance, suggesting that some features may be redundant within the existing feature set. (C) To further isolate the individual contribution of each feature to model performance, we performed single-metric ablation experiments (Fig. 4C), removing one feature at a time and evaluating its impact. The results indicated that removing F8 and F9 led to relatively notable improvements, with both AUC and accuracy increasing by more than 0.01. (D) Based on these findings, we constructed an optimized feature subset by excluding F8 and F9 and compared its performance against the baseline full feature set. The results (Fig. 4D) showed improvements, with AUC increasing by 0.032 and accuracy by 0.039. (E) Finally, we compared the performance of the full feature set and the optimized subset across observation windows from 1-10 minutes, and found consistent improvements in most windows, particularly in the 3-9 minute range (Fig. 4E).

**Fig. 4.**
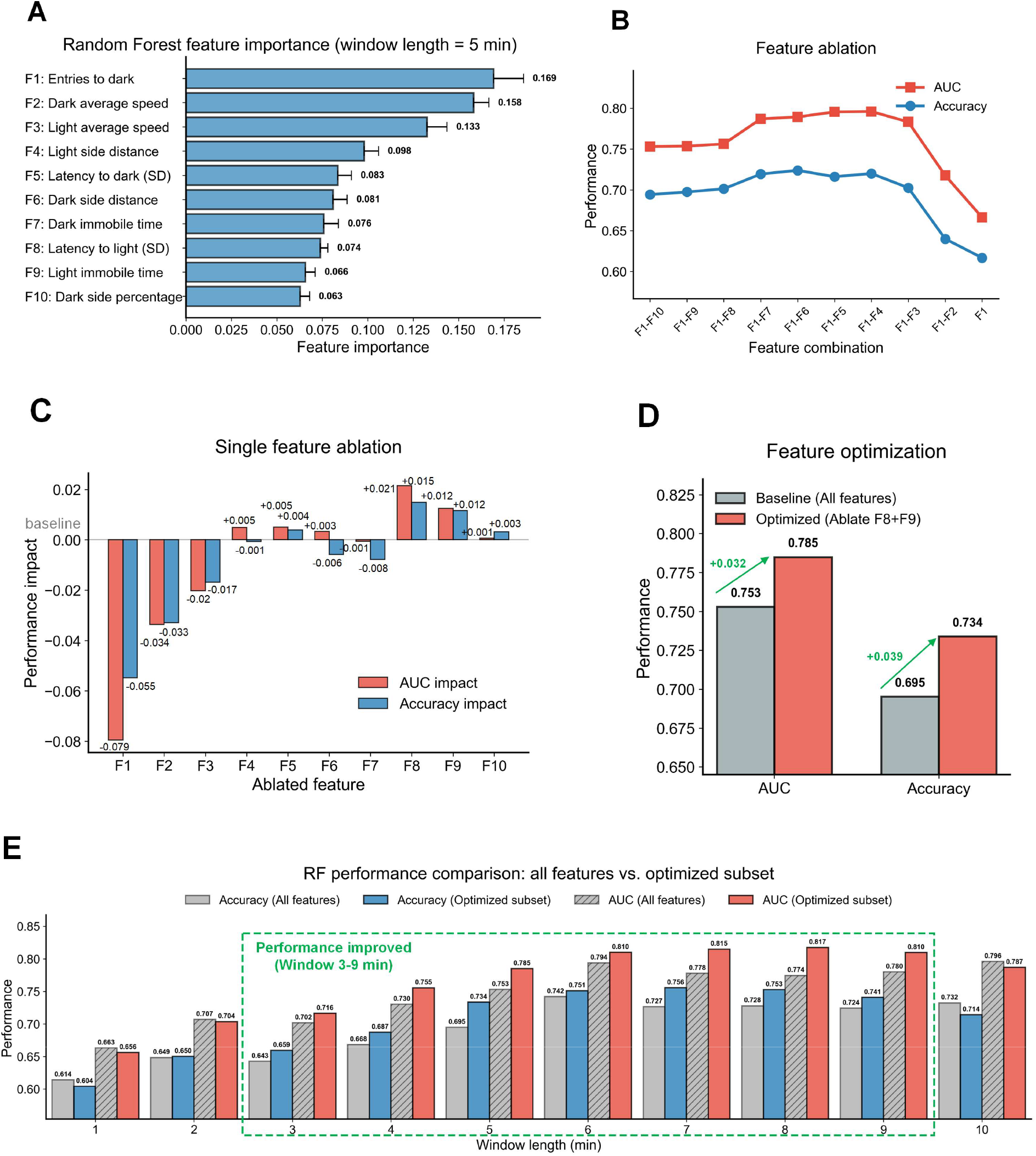
Feature importance analysis and performance improvement through feature subset optimization. **A**. Feature importance ranking of all ten behavioral features using the RF model. F1, F2, and F3 had the highest importance (weight > 0.1). **B**. Stepwise feature ablation based on inverse importance ranking showed that removing certain features (F9, F8, F7, F5) unexpectedly improved model performance. **C**. Single-feature ablation revealed that removing F8 and F9 led to the most notable performance gains. **D**. Comparison between the full feature set and the optimal subset (excluding F8 and F9) showed clear improvements (AUC improved by 0.032, accuracy by 0.039). **E**. Evaluation across observation windows from 1-10 minutes demonstrated that this optimization improved classification performance in most windows, particularly between 3-9 minutes.

It should be emphasized that the conclusions presented in Fig. 4 were derived from the mouse sample dataset used in this experiment, and their generalizability remains unknown. Therefore, the experiments in Fig. 4 are not intended to negate the role of certain behavioral features, but rather to demonstrate an approach for improving model performance through feature subset optimization. All experimental results reported earlier in this study were based on the full feature set, as this more comprehensively reflects the overall discriminative capacity of the model without relying on specific feature selection. In addition, although the feature importance analysis was conducted using the RF model, removing features F8 and F9 also led to performance improvements in the other four models, confirming the generalizability of this approach. For example, when the optimized feature subset was applied to the SVM model, the highest AUC improved from 0.836 to 0.866, and the highest accuracy increased from 0.784 to 0.802. The detailed results are provided in the Supporting Information.

## Discussion

In this study, we found that dwell time in the light or dark compartment, a commonly used single behavioral meatric in traditional light/dark box visual function assessments, showed clear limitations in distinguishing sighted from blind mice. In our experiments, the classification ability of the single-metric approach was close to random, and no significant separation was observed even when varying the observation window or habituation period settings. Interestingly, in the feature importance analysis of Fig. 4A, the percentage of dark-side dwell time was identified as the least important feature. As shown in Fig. 5, we summarized the dark side percentage distribution of the two groups of mice within a 20-minute experimental window. Among the 25 sighted mice, 52% (13 mice) clustered in the 40-50% range, and 32% (8 mice) exceeded 50%, with overall dark-side dwell percentages ranging between 30% and 70%. In contrast, the blind mice exhibited a more evenly distributed pattern with a broader range (20-80%), reflecting greater individual variability. However, previous studies were able to detect group-level differences using mean dark dwell time or percentage, despite considerable overlap in individual data. We speculate that the differences between our findings and previous reports may result from the following factors. First, different studies have employed different experimental settings, such as habituation duration and observation window length; our results suggest that these conditions indeed influence dark-side dwell time. Second, prior studies have shown that strain, age(or neuronal maturation) may also influence exploratory activity and anxiety-related behavior in the light/dark test, thereby affecting the interpretation of results^[8]^. In addition, our data showed that the blind group exhibited markedly greater behavioral variability (Fig. 1A and B, reflected by broader box plot distributions), suggesting that in the absence of visual drive, mouse behavior may be shaped by non-visual factors such as individual variability, anxiety, and exploratory motivation, thereby increasing group overlap. Our experiment suggest that it is possible that single dwell time is likely to subject to experimental conditions and non-visual factors, and therefore lacks stable discriminative power, although it is still not clear what all the possible confounding factors are. Given the same conditions, nevertheless, our data analyses suggest the value of a multi-feature approach. By integrating multiple behavioral features—including entries to dark, latency before crossing, locomotion speed, and immobility time, etc.—our method captures a more comprehensive behavioral profile, overcomes the limitations of single-metric assessment, and significantly improves the accuracy and robustness of visual function evaluation.

**Fig. 5.**
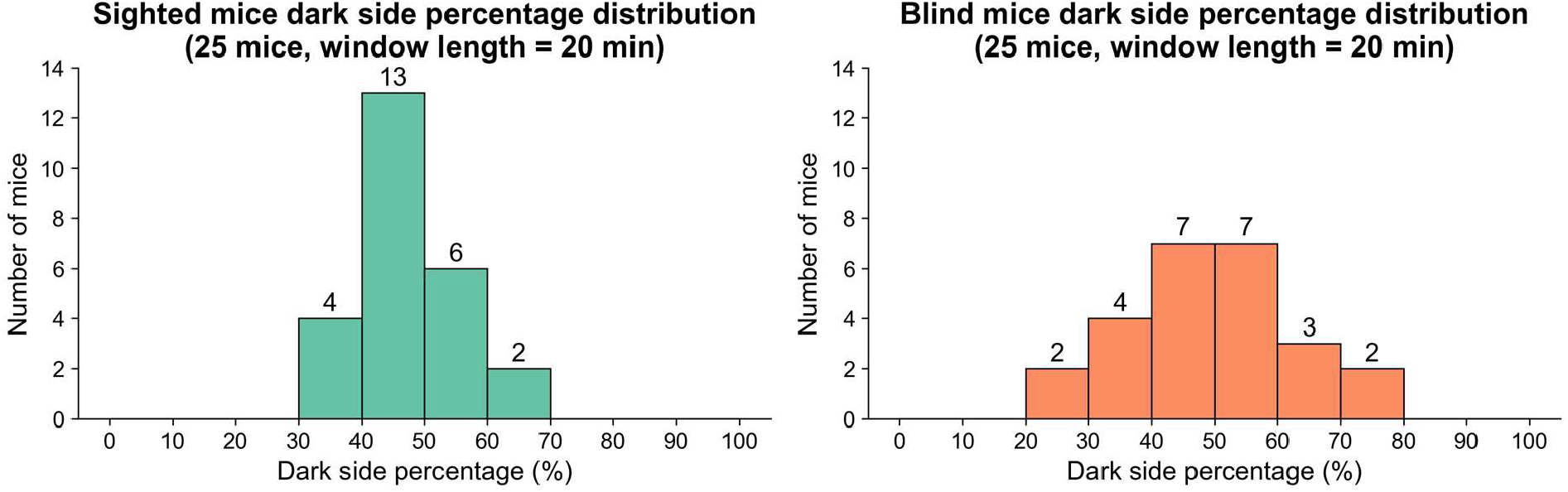
Distribution of dark side dwell percentages in sighted and blind mice over a 20-minute experimental window.

In terms of observation window design, this study set the maximum window length to 10 minutes, partly in reference to prior studies that commonly employed 3-10 minute settings^[9]- [16], [18]-[24]^, and partly because excessively long windows may reduce efficiency and hinder effective discrimination. Notably, our results showed that when the observation window length reached 2 minutes or longer, sighted and blind mice could be significantly distinguished (Fig. 1C), and when the window exceeded 5 minutes, the performance of the multi-feature machine learning model plateaus (Fig. 3A and B), indicating that reliable discrimination can be achieved without overly long observation periods. In addition, we adopted a sliding window strategy to segment the experimental data, which offered multiple advantages: first, it substantially increased the number of training samples, improving model training and helping to prevent overfitting due to limited data, thereby enhancing generalizability; second, it allowed coverage of the entire experimental session, capturing behavioral differences that may emerge at different time periods and avoiding the omission of critical information associated with a single window; and third, by functioning as a form of repeated sampling, the sliding window approach reduced the influence of random fluctuations in individual windows, making the overall results more robust and reliable.

Certain limitations of the current approach should be considered. First, the mouse strains and age ranges included in this study were limited; mice with different genetic backgrounds, sexes, or developmental stages may exhibit distinct behavioral patterns, and thus the generalizability of our results requires further validation. In addition, although we optimized behavioral features through feature importance analysis, feature engineering can still be improved, for example by incorporating additional potentially informative behavioral features—such as route distribution patterns, spatial occupancy distribution, or speed changes before and after crossings—or employing automated feature selection algorithms to further enhance model performance. Despite these limitations, the proposed multi-feature machine learning approach demonstrates clear advantages over conventional single-metric assessment. Once trained, the model enables automated and efficient visual function assessment, substantially improving accuracy and stability while reducing time and labor costs. These characteristics suggest that the method has the potential to serve as a more reliable tool for visual function assessment in preclinical studies. In the future, through standardized experimental protocols and software frameworks, this approach could be broadly applied to large-scale studies and cross-laboratory validation, providing enhanced technological support for visual function assessment and related disease research.

## Methods

### Animals

A total of 50 mice were tested, including 25 sighted and 25 blind mice. The sighted group included 8 wild-type (WT) mice aged 5-69 weeks and 17 young Rhodopsin knockout (*Rho*^−/−^) mice aged 4-10 weeks with preserved visual function. The blind group included 8 double optic nerve crushed *Rho*^−/−^ mice aged 10-24 weeks and 17 older *Rho*^−/−^ mice aged 16-45 weeks. *Rho*^−/−^ mice were generated at Trinity College in Dublin, Ireland (Humphries et al., 1997^[25]^) and were bred at the animal facility of the Schepens Eye Research Institute. The C57BL/6J WT mice were purchased from Jackson Laboratory (Bar Harbor, ME; Cat # 000664). Mice of both sexes were used in the experiments. *Rho*^−/−^ mice 6 weeks of age have normal vision with visual acuity (VA) of ∼0.45 cycle/degree, comparable to adult C57BL/6J wild-type mice. By 9 weeks, the VA in *Rho*^−/−^ mice decreases to ∼0.34 cycle/degree and continues to decline until complete blindness occurs by 4 months of age (Gunes et al., 2025^[26]^, and Xiao et al., 2019^[27]^). Thus, *Rho*^−/−^ mice were categorized as either sighted animals at 4-10 weeks of age, or blind animals beyond 16 weeks of age. Blind mice also included WT aged 2-5 months, which underwent optic nerve crush (ONC) surgery as described previously (Cho KS et al., 2005^[28]^; Tai WL et al., 2023^[29]^). Mice were housed in a temperature-regulated room with a 12-hour light/dark cycle and had free access to food and water. All animal experiments were approved by the Institutional Animal Care and Use Committees (IACUC) of the Schepens Eye Research Institute of Mass Eye and Ear and were conducted in accordance with the Association for Research in Vision and Ophthalmology (ARVO) Statement for the Use of Animals in Ophthalmologic and Vision Research.

### Automated light/dark box system

Fig. 6 shows the automated light/dark box system we built to assess light perception in rodents for different strains, ages, sexes and pathologies. The system is placed in a dark chamber and mainly composed of the following three units: (1) A light/dark box. The box consists of a light compartment and a dark compartment. The light and dark compartments measure 30 cm × 40 cm × 40 cm and 20 cm × 40 cm × 40 cm (length × width × height), respectively. An adjustable opening is located between the two compartments to allow rodents of different sizes to travel freely between them. A LED white light source (luminous flux: 180 lm, color temperature: 4000 K; corresponding to an estimated theoretical maximum illuminance of ∼1500 lux at the floor level of the light compartment, calculated as luminous flux divided by the floor area of 30 cm × 40 cm = 0.12 m^2^) and an infrared light source (wavelength: 940 nm, power: 12 W) are mounted on the top of the light and dark compartments, respectively, allowing the infrared camera to simultaneously monitor both areas. (2) An infrared camera mounted above the box captures real-time images of the animals at a rate of 30 frames per second (fps). (3) A computer (Intel i7-12700H CPU @ 2.30 GHz) records video data of the experiment and subsequently analyzes it using a self-developed rodent localization algorithm to continuously monitor the animal’s position and calculate various behavioral metrics. These metrics are then normalized and used as features for the machine learning model in both training and prediction tasks.

**Fig. 6.**
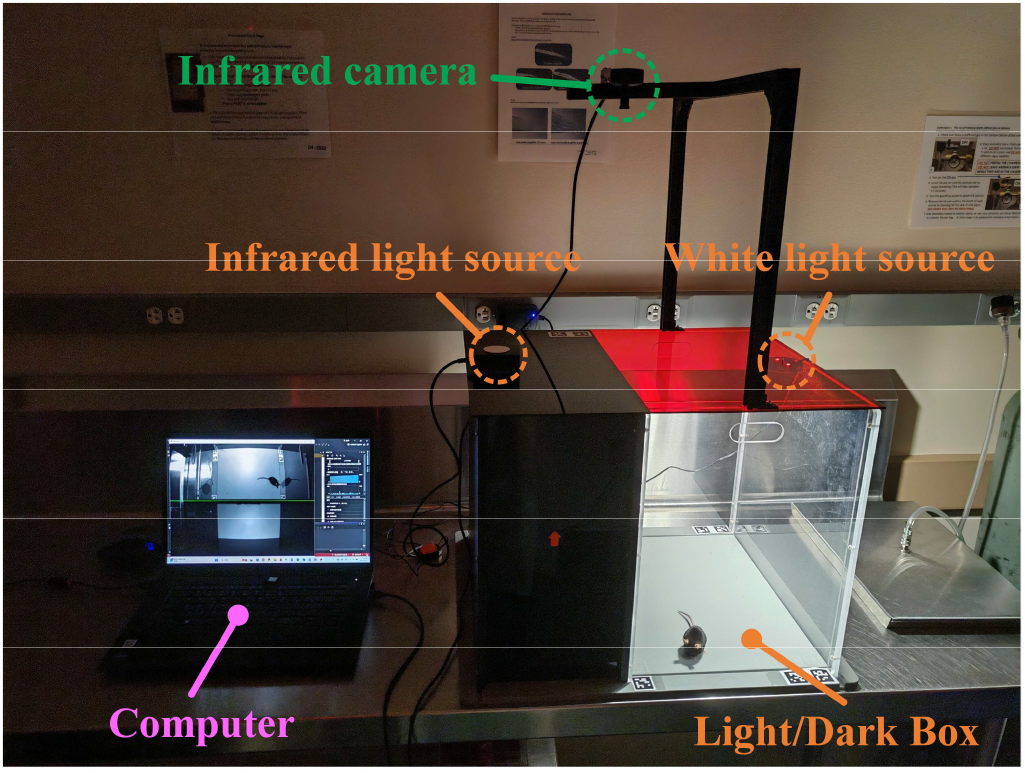
The automated light/dark box system.

### Rodent localization algorithm

To precisely localize the position of the test animal, we developed a rodent localization algorithm applicable to rodents such as mice and rats. As shown in Fig. 7, the algorithm mainly employs image processing techniques and consists of the following three steps. (1) Automatic region localization. Six fiducial markers^[30]^ were placed around the light compartment to assist in locating the regions corresponding to the light and dark compartments, as well as in correcting the camera angle. The program accurately detects the corner points of each marker to localize the light compartment area, and subsequently determines the dark compartment area based on the actual dimensional ratio between the two compartments. (2) Background subtraction was applied to separate the rodent from the static background. After adjusting the camera position, a background image was first captured. During the experiment, each incoming frame was compared pixel-wise with the background image, pixels with differences exceeding a predefined threshold were identified as foreground and labeled as white, while the remaining pixels were considered background and labeled as black. (3) Tail removal and localization. We observed that the shadow, curvature, and length of the tail could affect localization precision. Therefore, a distance transform was applied to compute, for each foreground pixel, its distance to the nearest background pixel. This distance map effectively highlights the thicker body region of the mouse, allowing thin tail structures to be identified and removed by thresholding pixels below a predefined distance value. As the edge regions of the main body were also partially eliminated, a dilation operation was subsequently performed to restore the body size. Finally, the rodent’s contour was extracted and the centroid was calculated to determine the position coordinates. The positional information of each frame was recorded for subsequent metric computation and analysis.

**Fig. 7.**
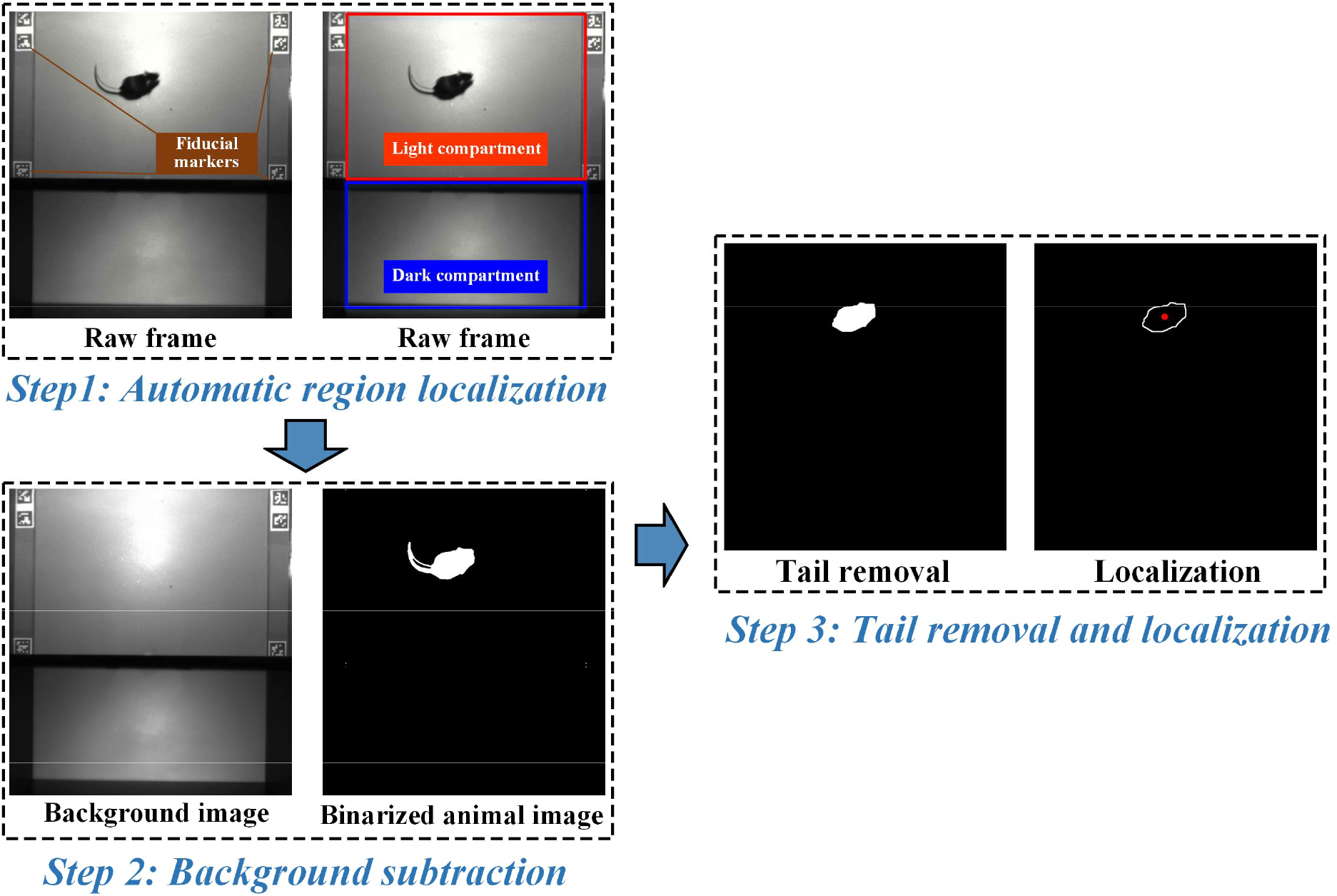
Workflow of the rodent localization algorithm. The algorithm consists of three main steps. **Step 1:** Automatic region localization. Six fiducial markers placed around the light compartment are detected to accurately define the light and dark compartments and correct camera angle. **Step 2:** Background subtraction. A background image captured before the experiment is compared with each incoming frame to extract the animal as a binarized foreground mask. **Step 3:** Tail removal and localization. Distance transform and thresholding are applied to remove thin tail structures, followed by dilation to restore body shape. The animal contour is then extracted, and the centroid of the body is calculated to obtain the position coordinates.

### Machine learning model

To construct a multi-feature machine learning paradigm, we first defined 10 behavioral metrics as input features for the machine learning models, as shown in Table I. Considering the inconsistency in feature scales and the potential nonlinear relationships among the selected features, which include parameters with different physical units and value ranges such as time, distance, and speed, all computed features were normalized before being input into the models. We selected five commonly used machine learning classifiers, as described below, for comparative analysis. All models were implemented using Python’s *scikit-learn* library, with XGBoost additionally utilizing the *XGBoost* library. To ensure reproducibility, the random seed was set to 0 in all model training procedures.

**Table I.** Mouse behavioral metrics.

| Metrics | Description | Unit |
| --- | --- | --- |
| Entries to dark | Number of entries into the dark compartment | count |
| Dark average speed | Average speed in the dark compartment | cm/s |
| Light average speed | Average speed in the light compartment | cm/s |
| Dark side distance | Total travel distance within the dark compartment | cm |
| Light side distance | Total travel distance within the light compartment | cm |
| Latency to dark (SD) | Standard deviation of time taken to enter the dark compartment from the light side | s |
| Latency to light (SD) | Standard deviation of time taken to enter the light compartment from the dark side | s |
| Dark immobile time | Duration of immobility in the dark compartment | s |
| Light immobile time | Duration of immobility in the light compartment | s |
| Dark side percentage | Dwell time percentage in the dark compartment | % |

Support vector machine (SVM)^[31]^: SVM achieves classification by finding the optimal separating hyperplane in a high-dimensional feature space, making it particularly suitable for classification tasks with small to medium-sized datasets. We employed an SVM model with a radial basis function (RBF) kernel, which effectively handles nonlinearly separable data. The regularization parameter *C* was set to 1.0, and the kernel coefficient *γ* was set to “scale.” Probability estimation was enabled to output classification probabilities.

Logistic regression (LR)^[32]^: As a classic linear classification method, logistic regression maps linear combinations to probability space through the logistic function. Although it is a linear model, logistic regression still demonstrates good performance in the classification tasks of this study after standardization preprocessing. The model was trained with L2 regularization and optimized using the LBFGS solver, with the regularization strength parameter *C* = 1.0. The maximum number of iterations was increased to 1000 to ensure convergence.

Multilayer perceptron (MLP)^[33]^: As a neural network method, MLP can learn complex nonlinear feature combinations and is suitable for handling classification problems with complex interactions between features. In this study, we employed a two-hidden-layer architecture (100 and 50 neurons) using the ReLU activation function and the Adam optimization algorithm, with an initial learning rate of 0.001 and a constant learning-rate schedule. The L2 regularization coefficient was set to 0.0001, and the maximum number of iterations was 500. To prevent overfitting, an early stopping mechanism was enabled with a validation fraction of 0.1, where training was terminated when the validation error showed no improvement for 10 consecutive iterations.

Random forest (RF)^[34]^: RF is based on an ensemble of decision trees, where each feature is independently evaluated for splitting, making it insensitive to feature scales and naturally suitable for inputs with mixed units. In this study, the number of trees was set to 100, and each tree was trained on randomly sampled subsets of features and data using the Gini impurity criterion. Bootstrap sampling was enabled by default.

Extreme gradient boosting (XGBoost)^[35]^: XGBoost is an ensemble learning method based on gradient boosting that improves overall performance by sequentially training weak learners and combining their prediction results. In this study, the model was configured with 100 estimators, a learning rate of 0.1, and a maximum tree depth of 6 under default settings. The evaluation metric was set to “logloss.”

### Data preparation and evaluation

We recorded 20-minute videos for each test mouse using the system shown in Fig. 6. The position of the mouse in each frame was then determined using the proposed animal localization algorithm. To construct training and testing samples, we defined an observation window of length *T*. For each video, the window was slid forward with a fixed interval *S* to generate multiple temporal segments. For each segment, the 10 behavioral metrics listed in Table I were computed. As a result, each video was divided into multiple samples, and the extracted metrics were normalized and used as feature inputs for model training and testing. We employed K-fold cross-validation with group-wise partitioning to evaluate model performance. The group-wise partitioning ensured that data from the same mouse did not appear in both training and testing sets simultaneously. In this approach, mice were treated as the unit of grouping, and all individuals were evenly divided into *K* folds, ensuring that each fold contained the same number of normal-vision and blind mice. For each training process, the model was re-initialized, and one fold was used as the test set while the remaining folds served as the training set. In this study, we used 5-fold cross-validation (*K*=5), resulting in five trained models, with each fold containing 5 sighted mice and 5 blind mice to maintain class balance. After completing all *K* rounds, the test predictions from all folds were aggregated to cover the entire dataset, and the overall AUC and accuracy were then computed based on these combined predictions. This cross-validation strategy helps ensure that the evaluation is robust and less sensitive to how the data is split, providing a reliable estimate of the model’s generalization performance.

## Supporting Information

**S1 Fig.**
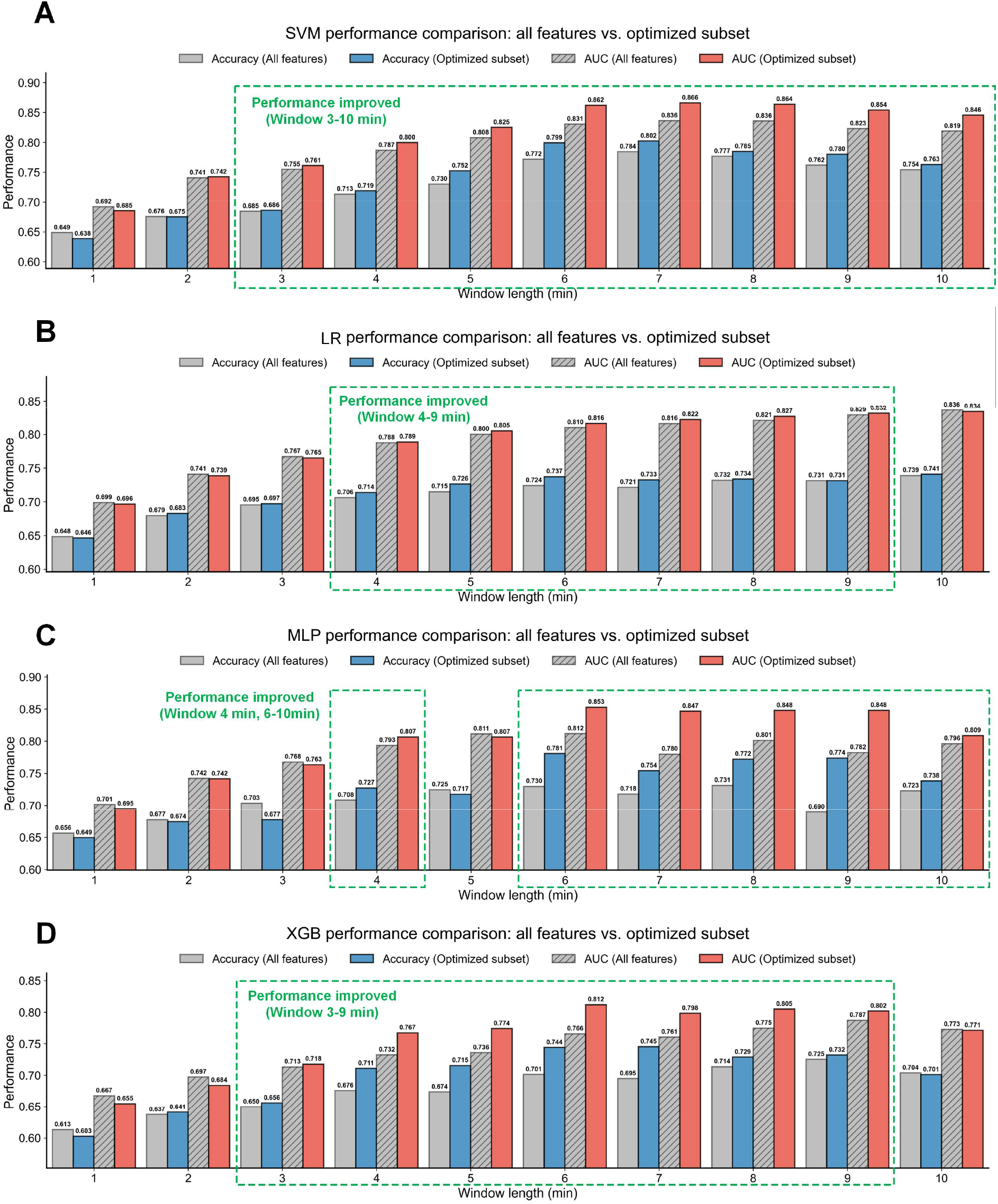
Performance comparison of different models using full feature set versus optimized feature subset. **A**. SVM: Improvements observed across 3-10 minute windows, with maximum accuracy increasing from 0.784 to 0.802 and AUC from 0.836 to 0.866. **B**. LR: Slight improvements in the 4-9 minute range, with maximum accuracy increasing from 0.724 to 0.737 and AUC from 0.829 to 0.832. **C**. MLP: Performance enhancement in 4-minute and 6-10 minute intervals, with maximum accuracy increasing from 0.730 to 0.781 and AUC from 0.812 to 0.853. **D**. XGB: Consistent improvements across 3-9 minute intervals, with maximum accuracy increasing from 0.695 to 0.745 and AUC from 0.766 to 0.812.

## Acknowledgements

This research was supported by the Advanced Research Projects Agency for Health (ARPA-H OTA #1AY2AX000056) in part. The views and conclusions contained in this document are those of the authors and should not be interpreted as representing the official policies, either expressed or implied, of the United States Government.

## Author Contributions

**Conceptualization:** Gang Luo, Dong Feng Chen.

**Data curation:** Tengxiao Wang, Chen-Yuan Lee.

**Formal analysis:** Tengxiao Wang, Gang Luo.

**Funding acquisition:** Gang Luo.

**Investigation:** Tengxiao Wang, Chen-Yuan Lee, Karen Chang, Gang Luo.

**Methodology:** Tengxiao Wang, Gang Luo.

**Project administration:** Gang Luo, Dong Feng Chen.

**Resources:** Karen Chang, Matteo Tomasi, Dong Feng Chen.

**Software:** Tengxiao Wang.

**Supervision:** Gang Luo, Dong Feng Chen.

**Validation:** Tengxiao Wang, Chen-Yuan Lee.

**Visualization:** Tengxiao Wang.

**Writing – original draft:** Tengxiao Wang, Karen Chang, Gang Luo.

**Writing – review & editing:** Tengxiao Wang, Gang Luo.

## Data Availability Statement

All raw data and analysis code supporting the findings of this study will be made publicly available on GitHub or Zenodo upon publication.

